# Utilising multi-modal data-driven network analysis to identify monotherapy and combinational therapy targets in SOX2-dependent squamous cell lung cancer

**DOI:** 10.1101/2025.01.30.635634

**Authors:** Woochang Hwang, Daniel Kottmann, Wenrui Guo, Méabh MacMahon, Lucia Correia, Rebecca Harris, Frank McCaughan, Namshik Han

## Abstract

Drug discovery requires a deep understanding of disease mechanisms, making the integration and analysis of diverse, multi-modal data types essential. These data types, spanning omics types, disease-associated, and biological pathway information, often originate from disparate sources and must be combined to uncover critical insights. We have developed iPANDDA (*in-silico Pipeline for Agnostic Network-based Drug Discovery Analysis*), a computational pipeline that integrates multi-modal data through a network-based approach to predict candidate drug target proteins for specific diseases.

We applied iPANDDA to lung squamous cell carcinoma (LUSC), a subtype of non-small cell lung cancer (NSCLC) that accounts for approximately 25% of all lung cancer cases globally. Despite advances in cancer therapeutics, targeted therapies specifically approved for LUSC remain lacking, exacerbated by the shortage of robust models for studying LUSC carcinogenesis and therapeutic responses. The SOX2 gene, amplified in about 50% of LUSC patients, plays a critical role in driving and maintaining the cancer phenotype. Using iPANDDA, we identified relevant therapeutic targets for SOX2-dependent LUSC.

Selected candidate drug targets were validated in vitro using cell-based models. We conducted target inhibition studies in both SOX2-dependent and non-SOX2-dependent cell lines, evaluating the effects of inhibition and knockout through cell viability assays.

Our findings confirmed key monotherapy and combination therapy targets for SOX2-focused models. Specifically, we validated the AKT and mTOR complexes as promising therapeutic targets for LUSC. Additionally, we identified potential pathways for developing novel combination therapies targeting SOX2-dependent LUSC. iPANDDA offers a robust approach to refining and focusing therapeutic strategies for diseases with unmet clinical needs.

## Introduction

Drug discovery is a lengthy, costly process with a high failure rate, often due to insufficient efficacy in the target disease^1^. This failure frequently stems from a limited understanding of disease pathogenesis and the role of specific drug targets in the disease. The increasing availability of datasets on individual diseases and potential drug targets has spurred efforts to integrate disparate biological data to enhance target identification and streamline drug development^1,2^. Additionally, combination therapies are gaining attention for their potential to improve therapeutic outcomes^3–5^. However, integrating diverse datasets to predict effective monotherapies and combination therapies remains a significant challenge, highlighting the unmet need for computational algorithms capable of prioritizing therapeutic targets in a disease-specific context.

Lung cancer is the leading cause of cancer-related deaths worldwide. Lung squamous cell carcinoma (LUSC), a subtype of non-small cell lung cancer (NSCLC), accounts for approximately 25% of all lung cancer cases^6^. Despite advances in cancer therapeutics, no targeted therapies have been approved for LUSC. Among its key molecular drivers, amplification of the 3q26 locus is observed in about 50% of LUSC patients^7–10^. Evidence suggests that SOX2 is a critical target gene of this amplification and a key molecular driver of LUSC pathogenesis. However, targeting transcription factor s (TFs) like SOX2 poses significant therapeutic challenges due to their promiscuous interactions with other proteins and DNA binding sites. These complexities have hindered efforts to define the critical upstream and downstream proteins or pathways influenced by SOX2, resulting in the absence of targeted treatments for this devastating disease.

Recent advances in molecular profiling have generated a wealth of publicly available ‘omics’ datasets for LUSC^9^, along with functional datasets from CRISPR and RNAi screens^11^. However, integrating these complex and heterogeneous datasets to derive actionable insights remains a formidable task. Sophisticated analytical tools that integrate multi-modal datasets are urgently needed to comprehensively understand the pathobiology of LUSC and identify novel therapeutic strategies.

To address these challenges, we developed iPANDDA (*in-silico Pipeline for Agnostic Network-based Drug Discovery Analysis*), a computational algorithm designed to integrate and analyze multi- modal datasets. Using iPANDDA, we constructed a meaningful protein-protein interaction (PPI) network centered on SOX2 and LUSC, identifying therapeutically vulnerable nodes. These targets were then ranked based on disease relevance and druggability.

This systematic approach identified candidate targets for SOX2-dependent LUSC, which we validated experimentally. iPANDDA successful identified known therapeutic targets, thus validating our approach, as well as identifying a number of promising novel targets and drug candidates for this unmet clinical need.

## Results

### iPANDDA: a pipeline to identify candidate drugs and druggable pathways for diseases associated with specific genes of interest

iPANDDA is a pipeline developed to identify candidate drugs and druggable pathways for diseases with a specific genetic dependency but for which treatment options are limited. We developed iPANDDA using an agnostic data-driven approach to construct a network-based model of the disease of interest with the aim of identifying key disease nodes and potential drug targets. The input of the iPANDDA pipeline is multi-modal data related to the disease; the outputs are candidate drugs and targets or networks that should be a focus of therapeutic efforts.

In this report, the focus is on a subset of LUSC that is SOX2-dependent and for which there are no specific therapeutics available. The pipeline is summarised in Figure 1A. There are five key computational steps prior to wet lab validation experiments. These are summarised below, with further details provided in the Methods section. The novelty of iPANDDA lies in the breadth of data feeding into the algorithm, its integration, and the incorporation of experimental data specifically to disease being examined, in this case SOX2-dependent LUSC.

1. **Multi-Modal Datasets Related to SOX2-Dependent LUSC:** iPANDDA utilises proprietary and publicly available multi-modal data associated with LUSC, comparative data from normal tissue, and additional proprietary experimental data.

- **Proprietary data** are derived from an in vitro organotypic model system in which the driver oncogene SOX2 is acutely deregulated in an immortalised airway epithelial cell line. This model aims to recapitulate the molecular, cellular, and physiological environment in which LUSC develops.
- **Public datasets** include cancer-specific genomic and transcriptomic data from The Cancer Genome Atlas (TCGA)^12^, normal tissue genomic and transcriptomic data from the Genotype-Tissue Expression (GTEx) database^13^, and data from the Open Targets and OmniPath databases^14^. Open Targets focuses on disease-target associations, while OmniPath is focused on cell signalling networks. Additionally, data from the ENCODE database^15^ on transcription factor (TF) activity and binding sites for SOX2 are incorporated.
2. **Data Integration and Network Construction:** This step integrates all the information gathered from the multi-modal disease association study (Step 1) into a single data space, forming a network (graph-based data structure). The network includes PPIs of genes of interest, disease-associated genes from Open Targets, known drug targets, TFs for genes of interest, and differentially expressed genes (DEGs) from integrated transcriptomics data. This integrated network serves as the foundation for constructing a disease-specific network that encapsulates the entire disease signature.
3. **Network Analysis:** The network is analysed to identify key proteins that are central to the SOX2 network and, therefore, may represent ideal therapeutic targets. Key proteins are determined using multiple network centrality algorithms and the Random Walk with Restart (RWR) algorithm. CRISPRi data from the Dependency Map (DepMap)^16^ is used to further refine the network to retain only those with demonstrable LUSC dependency.
4. **In-Silico Drug Simulation:** This step stratifies potential candidate drugs using a network proximity method^17^ to predict how closely drug targets correspond to the identified key proteins in the network.

- **Druggable targets** are those for which an existing compound to target^18^ them exists based on the proximity analysis.
- **Undruggable targets** are those for which no existing compound is available, as suggested by the drug proximity analysis.
5. **Target Prioritisation:** Using iPANDDA, we rank druggable and undruggable targets according to a stepwise network analysis score. Prioritisation is based on a combination of factors, including the network analysis score, druggability assessment, structural clustering, RNA expression levels, and the results from the in-silico drug simulation. One of the advantages of creating a network is that it may suggest novel combinations of therapeutics targeting separate nodes within the network. Combination therapeutics are prioritised based on their potential for targeting more than one functional pathway within the network.
6. **In Vitro Validation:** We evaluate the impact of therapeutic compounds on SOX2-dependent and non-SOX2-dependent cell lines using a range of concentrations, analysing cell viability with a resazurin-based assay, and presenting results as dose-response curves.

**Figure 1.**
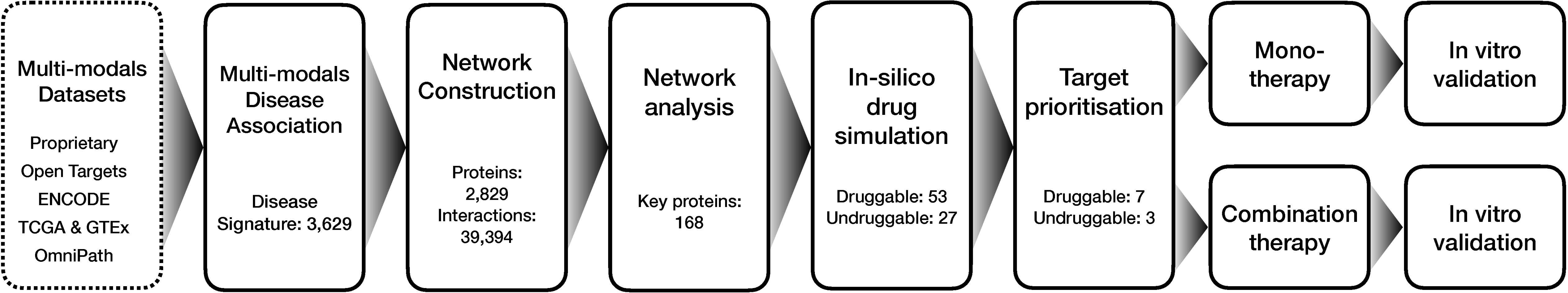
Overview of iPANDDA. iPANDDA is a computational pipeline designed to identify candidate drugs and therapeutic pathways for diseases of interest. For SOX2-dependent LUSC, multi-modal datasets were integrated to construct a disease-specific protein-protein interaction (PPI) network. Data sources included transcriptional signatures (1,636 genes identified from LUSC-specific TCGA analysis, 2,312 DEGs from a bespoke SOX2-dependent model using RNA-seq and ChIP-seq), transcription factors (12 from ENCODE), LUSC dependency data (157 from Open Targets), and SOX2-interacting proteins (12 from OmniPath). Network analysis using centrality algorithms and Random-Walk identified 168 core proteins. Drug simulation was applied to these core proteins to classify them as druggable or undruggable. Candidate drug-targets were prioritised using a combination of network and omics analyses, leading to in vitro validation experiments for 7 monotherapy candidates and the identification of novel combination therapy strategies.

### Identification of a multi-modal disease signature for constructing SOX2-dependent LUSC network

We interrogated multiple datasets associated with SOX2-dependent LUSC to establish a multidimensional disease-specific signature.

#### LUSC/SOX2 Transcriptional and Functional Signature

We utilized proprietary RNA-seq datasets and the TCGA database to identify transcriptional signatures specific to SOX2-dependent LUSC. In the proprietary RNA-seq dataset, SOX2 was confirmed as the most highly upregulated gene, and DEGs were identified (Figure 2A). To further analyze the regulatory role of SOX2 in LUSC, we incorporated SOX2 ChIP-seq datasets, which revealed direct binding sites and provided insights into its transcriptional regulatory network (Figure 2B).

**Figure 2.**
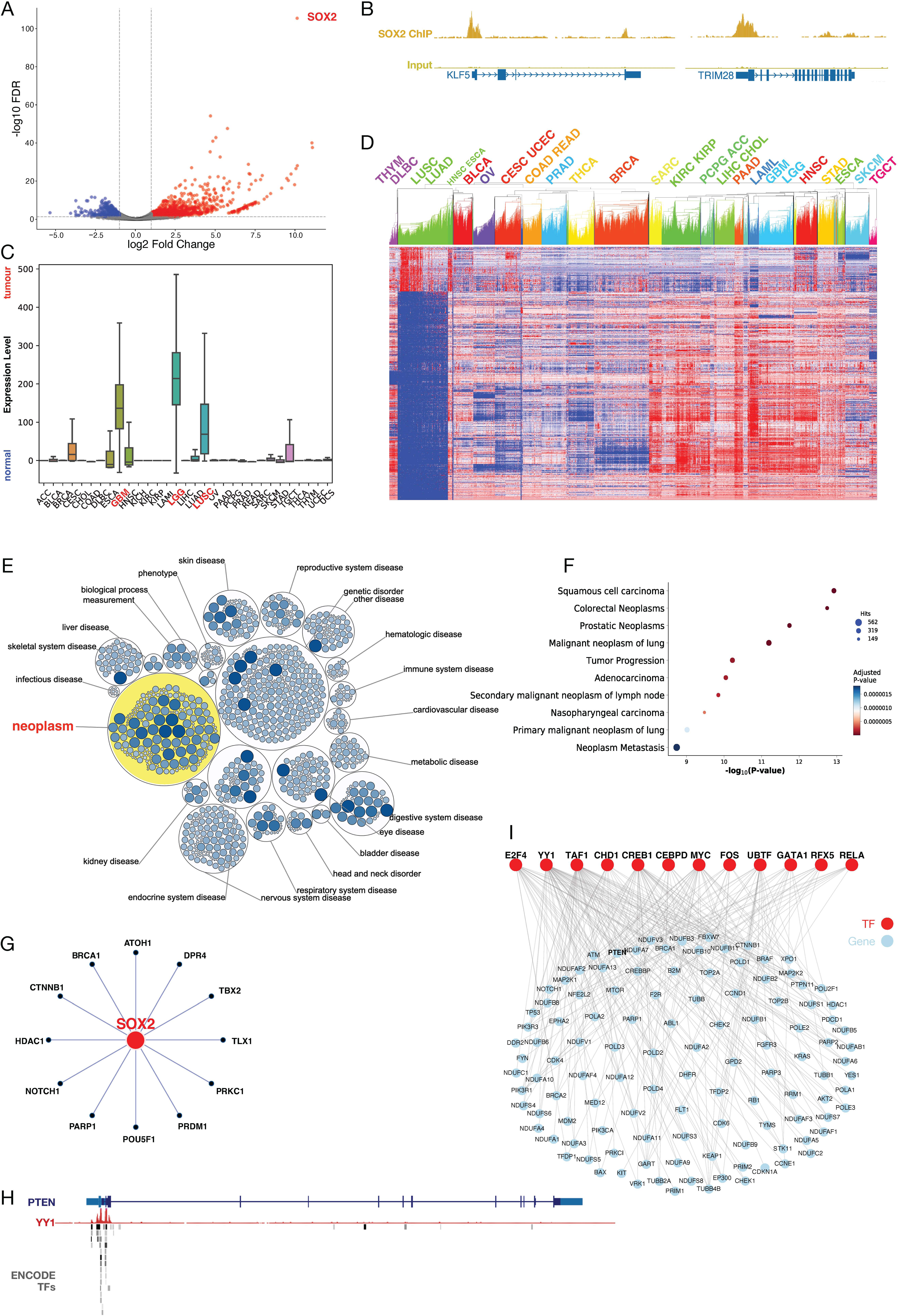
Multi-modal data integration and disease signatures. **A)** Differential expression analysis from proprietary RNA-seq data identified SOX2 as the most upregulated gene in SOX2-dependent LUSC. **B)** SOX2 ChIP-seq analysis revealed direct binding sites of KLF5 and TRIM28, providing insights into its transcriptional regulatory role. **C)** Pan-cancer analysis of SOX2 expression in normal versus cancer samples using ENCODE and TCGA data confirmed significant SOX2 upregulation in LUSC. **D)** Heatmap of TCGA samples clustered based on LUSC-specific cancer genes identified through comparative analysis of LUSC versus other cancer types. Rows represent individual LUSC-specific genes, and columns represent 31 TCGA cancer types. Red indicates upregulation, and blue indicates downregulation. Lung cancer types (LUSC and LUAD) cluster together, as do other foregut-origin cancers, including squamous types (LUSC, HNSC, and ESCA). **E)** Disease enrichment analysis using OmniPath confirms SOX2’s predominant association with neoplasms. **F)** DisGeNet over-representation analysis demonstrates that SOX2-dependent DEGs are significantly enriched in epithelial carcinomas, particularly squamous subtypes. **G)** Protein-protein interaction (PPI) network from OmniPath shows 12 direct interactors of SOX2. **H)** Example of transcription factor binding sites (TFBS) from ENCODE, showing YY1 binding to the PTEN gene. **I)** Regulatory map of 12 transcription factors upstream of SOX2, highlighting their interactions with downstream target genes in SOX2-dependent LUSC.

From the TCGA database, we conducted a cross-comparison of LUSC with the remaining 30 cancer types in the pan-cancer dataset to identify distinctly differentially expressed genes in LUSC. To enhance robustness, we integrated normal samples from TCGA and GTEx databases, overcoming the limitations of insufficient normal samples in the TCGA dataset (Supplementary Figure 1A). Instead of raw expression values, we used fold-change values to better capture differential expression. SOX2 showed significantly upregulated expression in LUSC compared to normal tissue (Figure 2C). Using an ANOVA test (p-value ≤ 1e-150), we identified 1,636 genes with enriched differential expression in LUSC (Supplementary Table 1). Hierarchical clustering of the pan-cancer TCGA dataset using these 1,636 genes showed that LUSC could be readily distinguished from other tumor types, with the nearest clusters comprising foregut-origin cancers, including lung adenocarcinoma (LUAD), head and neck squamous cell carcinoma (HNSC), and esophageal carcinoma (ESCA) (Figure 2D). Further validation using the Human Protein Atlas enrichment test demonstrated that these genes are significantly associated with lung-specific biology (Supplementary Table 2).

Using the Open Targets database, we confirmed that SOX2 is predominantly associated with neoplasms (Figure 2E). Additionally, we incorporated 62 LUSC-associated genes with an association score ≥0.8. This threshold was optimized using disease enrichment tests across gene lists, yielding the lowest p-value (6.0E-34) for LUSC. Furthermore, target proteins for drugs in Phase 3 or higher clinical trials for LUSC were also retrieved from the Open Targets database for inclusion in the analysis.

#### SOX2 Dependency Signature

Next, we focused on SOX2 dependency by analyzing DEGs from a proprietary organotypic model (OTC) and publicly available datasets, as well as interrogating existing datasets for the SOX2 interactome or predicted dependency. In the OTC model, SOX2 deregulation in immortalized human bronchial epithelial cells identified 2,312 DEGs (|logFC| ≥ 1, FDR < 0.05). These genes were significantly enriched in epithelial carcinomas, particularly squamous subtypes, as shown by the DisGeNet^19^ over-representation protocol (Figure 2F).

We then explored proteins directly interacting with SOX2 or serving as putative upstream transcriptional regulators of SOX2. From the Omnipath database, we identified 12 proteins with direct protein-protein interactions (PPIs) involving SOX2 (Figure 2G). To identify upstream transcriptional regulators of SOX2, we used the ENCODE Transcription Factor Binding Sites (TFBS) database. A transcription factor (TF) over-representation analysis (p-value < 0.01) identified 12 TFs, including YY1, MYC, and CREB1. Figure 2H illustrates an example of TFBS mapped to the PTEN gene, highlighting how these regulators influence key downstream targets. The regulatory map of the identified 12 TFs and their associated target genes is shown in Figure 2I, emphasizing the extensive transcriptional control exerted by these TFs in SOX2-dependent LUSC.

#### Construction of LUSC SOX2-dependent (LUSOX) network

To construct a comprehensive SOX2-dependency network for LUSC, we integrated disparate datasets encompassing transcriptional, functional, and interaction profiles (Supplementary Table 1). This network serves as a foundation to unravel SOX2’s multifaceted role in LUSC pathogenesis and to identify potential therapeutic targets. Using the STRING database^20^, we developed a protein-protein interaction (PPI) network comprising 2,829 proteins and 39,394 interactions. Within this network, SOX2 directly interacts with 156 proteins. These directly interacting proteins connect further with their first-order neighbours, forming a broader network of interactions. These neighbour proteins also interact with LUSC-associated genes, culminating in a network that includes 1,234 genes, approximately half of the entire network. This extensive connectivity demonstrates SOX2’s significant influence on the network through direct interactions, neighbour proteins, and connections to LUSC-associated genes. These findings underscore SOX2’s central role in LUSC pathogenesis and its potential as a key target for therapeutic intervention.

### Unveiling of LUSC disease mechanisms by network community detection

To elucidate the influence of SOX2 on LUSC disease mechanisms, we used a network community detection method (Louvain protocol – see Methods) to group the LUSOX network into eight distinct communities, each related to specific pathways (Figure 3A). These communities correspond to key biological processes such as RHO GTPase signalling, the Wnt pathway, keratinization, cytokine signalling, transcription pathways, interferon signalling, the cell cycle, and lipid metabolism, all of which are implicated in LUSC pathogenesis^21,22^.

**Figure 3.**
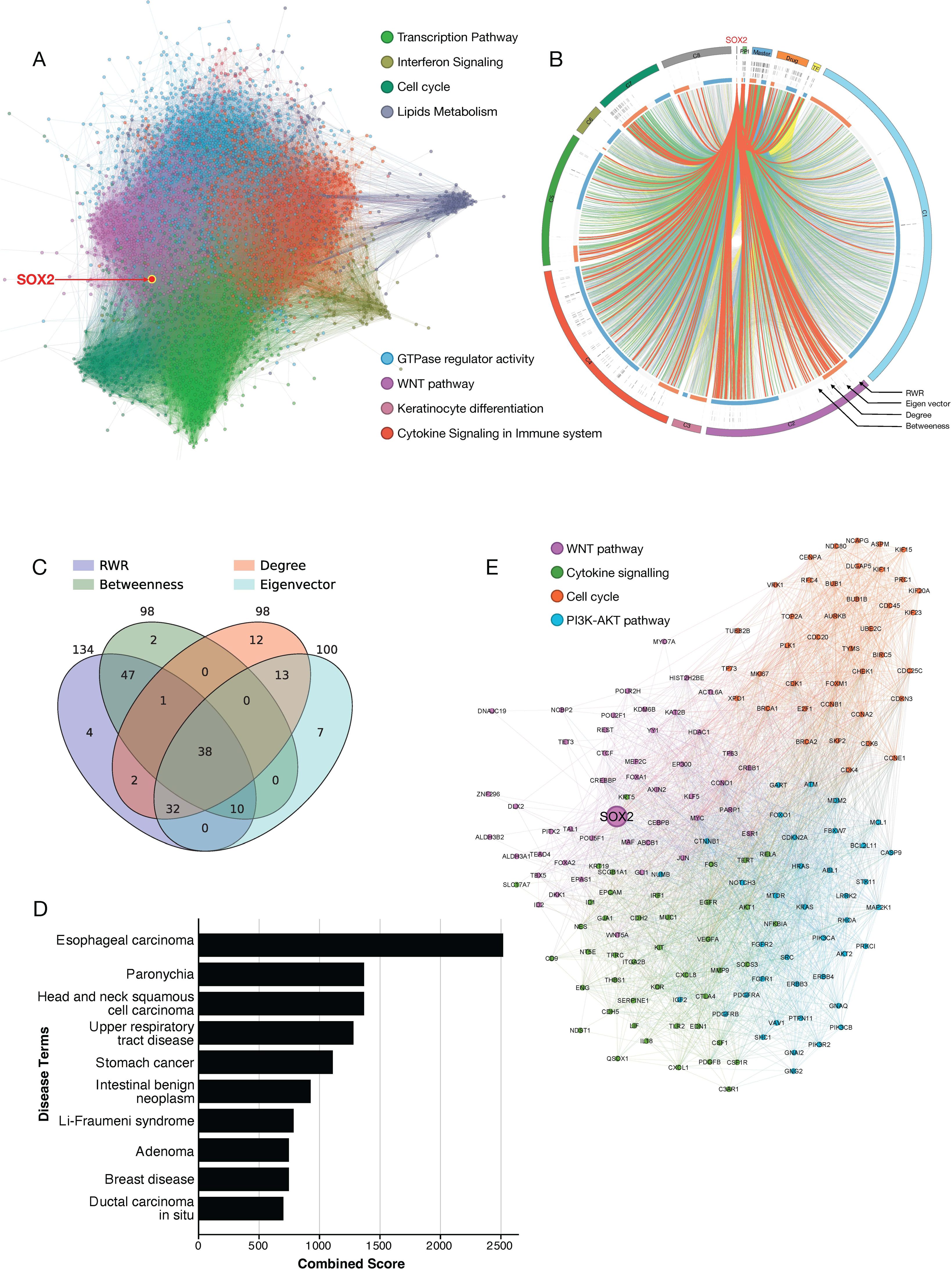
Network-Based Analysis of the SOX2-Dependent LUSC Pathways. **A)** The SOX2-dependent network consists of 2,829 genes and 39,394 interactions. The network is divided into eight communities, each associated with key pathways, including the Cell Cycle, Cytokine Signaling, Interferon Signaling, Lipid Metabolism, RHO GTPase Cycle, and WNT Pathway. Transcription Pathway refers to the community where key transcription factors and their target genes cluster, reflecting their central regulatory role. **B)** Circos plot of the SOX2-dependent network showing protein interactions (inner links), fold change values in LUSC (outermost track: red indicates upregulation, blue indicates downregulation), and network centrality scores (RWR, Eigenvector, Degree, and Betweenness centralities). The outer band colors represent pathway communities, consistent with panel A. SOX2 interacts directly with master regulators of LUSC, including transcription factors and proteins in cytokine signaling and WNT pathways. These master regulators bridge SOX2 to the broader network of 2,829 proteins, underscoring SOX2’s influence across pathways. Proteins associated with LUSC drugs represent targets from clinical or preclinical studies for lung squamous cell carcinoma, and the cytokine signaling pathway contains the highest number of core proteins. **C)** Venn diagram showing the overlap of core proteins identified through network analysis methods: Random Walk with Restart (RWR), Degree centrality, Betweenness centrality, and Eigenvector centrality. A total of 143 proteins (85%) were identified by more than one method, highlighting their critical role in the SOX2-dependent network. **D)** Over-representation analysis (ORA) of diseases associated with key proteins in the network, identifying squamous cell carcinoma as the most enriched disease. **E)** The core protein network is grouped into four communities, each representing distinct pathways: Cell Cycle, PI3K-AKT Pathway, Cytokine Signaling, and WNT Pathway/DNA Repair. The community containing SOX2 is strongly associated with the WNT Pathway and DNA Repair processes, emphasizing SOX2’s central regulatory role in oncogenic pathways.

We visualised these interdependencies using a Circos plot (Figure 3B, Supplementary Figure 2), which highlights the intricate network of biological pathways influenced by SOX2. The plot illustrates interactions between SOX2 and its directly interacting proteins (PPI), LUSC master regulators (Master), target proteins of LUSC drugs (Drug), transcription factors (TF) of LUSC signature genes, and differentially expressed genes in key pathways. These key pathways were labelled C1 through C8, with each representing a unique functional role: RHO GTPase signalling (C1), the WNT pathway (C2), keratinization (C3), cytokine signalling in the immune system (C4), transcription pathways (C5), interferon signalling (C6), the cell cycle (C7), and lipid metabolism (C8). SOX2 not only directly interacts with key proteins within these pathways but also exerts indirect control through the “Masters,” “Drug Targets,” and TFs. The arcs connecting the clusters (C1 to C8) emphasize SOX2’s pleiotropic role across multiple pathways implicated in LUSC, consistent with preclinical data suggesting it acts as a master “switch” for LUSC^21–23^.

These pathways include critical processes such as the Wnt pathway, which has a complex, context-dependent regulatory relationship with SOX2 and is strongly implicated in development and carcinogenesis. Similarly, RHO GTPases are key signalling molecules associated with cancer hallmarks, while deregulation of the cell cycle is a defining feature of cancer. Keratinization is a hallmark histopathology feature of LUSC. The network also implicated key immunological pathways, including cytokine signalling and interferon signalling, which could shape the tumour microenvironment (Supplementary Figure 2). This comprehensive network analysis confirms that SOX2’s influence spans multiple interconnected routes, potentially opening new avenues for therapeutic intervention.

### Discovery of drug targets for SOX2-dependent LUSC by network analysis

To further refine the best targets in SOX2-dependent LUSC, we applied multiple network analysis algorithms to the LUSOX network, including eigenvector centrality, degree centrality, betweenness centrality, and random walk with restart (RWR). Each algorithm provides unique insights into network dynamics. Eigenvector centrality identifies the most influential proteins by considering both the number of interactions and the importance of interacting proteins. Betweenness centrality identifies proteins that act as bridges between biological pathways. RWR quantifies how information spreads from a target protein to all other proteins in the network, detecting proteins most influenced by the target.

We performed 1,000 permutation tests for each algorithm to identify significantly associated proteins with empirical p-values less than 0.05 (see Methods). Using these methods, we identified 168 key proteins (Figure 3C, Supplementary Table 2). Figure 3C provides a Venn diagram showing the overlap among the proteins identified by the four-centrality metrics. Notably, 143 proteins (85%) were identified by at least two centrality algorithms, demonstrating strong agreement among the methods. Furthermore, 38 proteins were shared across all four algorithms, highlighting their centrality and potential as key targets in the LUSOX network.

Over-representation analysis (ORA) using the DISEASE database revealed that esophageal carcinoma, which is SOX2-associated squamous carcinoma, was the top enriched disease for these key proteins (Figure 3D). Figure 3D visualizes the results of the ORA, linking the identified proteins to squamous carcinoma and other related foregut-origin diseases, including esophageal carcinoma and head and neck squamous cell carcinoma. These diseases are also closely related to LUSC in the TCGA cluster (Figure 2D), further validating the biological relevance of the identified targets. Additionally, the PI3K-AKT pathway, the most altered pathway in LUSC, emerged as the top enriched biological pathway across all centrality algorithms^24^.

To further explore the biological implications, we constructed core networks from the 168 key proteins to investigate their functions and the biological pathways within the core network. Utilizing community detection algorithms, we introduced a novel approach for pathway identification that diverges from traditional gene enrichment tests. This unbiased method systematically identifies protein communities, corresponding to biological pathways, within a mathematical framework. By reducing reliance on potentially biased pathway databases, this approach offers a more objective means of pathway detection.

Through these community detection algorithms, we identified four distinct communities (Figure 3E), each associated with specific biological functions: the WNT pathway, PI3K-AKT pathway, cytokine signalling, and cell cycle regulation. Figure 3E illustrates the core network, with the four identified pathways distinctly colour-coded to highlight their respective clusters. For example, the WNT pathway is shown in purple, cytokine signalling in green, the cell cycle in orange, and the PI3K-AKT pathway in blue. SOX2’s central position within the network underscores its pleiotropic influence across these interconnected pathways.

This comprehensive map of interactions and pathways provides a critical resource for identifying and developing treatments that target distinct regulatory mechanisms within the disease.

### Network-based druggable target prioritisation and scoring

To prioritise therapeutic targets for SOX2-dependent LUSC, we utilised 168 key proteins identified through network analysis in the previous section. These key proteins formed the basis for drug simulations, where we analysed their potential to interact with 6,009 compounds from DrugBank^18^, encompassing approved, investigational, and experimental drugs. The simulations identified 521 drugs as potential candidates targeting the key proteins. Figure 4A visualises the focused PPI network of these targeted proteins, with node sizes reflecting the number of drugs targeting each protein. We then analysed the subset of 2,626 drugs with known indications and identified 217 drugs as potential LUSC therapeutics. Among these, 114 of the predicted drugs overlapped with cancer drugs (of the 2,626 drugs, 361 were classified as cancer drugs) (Supplementary Table 2). This overlap was statistically significant (hypergeometric p-value = 2.38E-27), supporting the relevance of the identified drug candidates.

**Figure 4.**
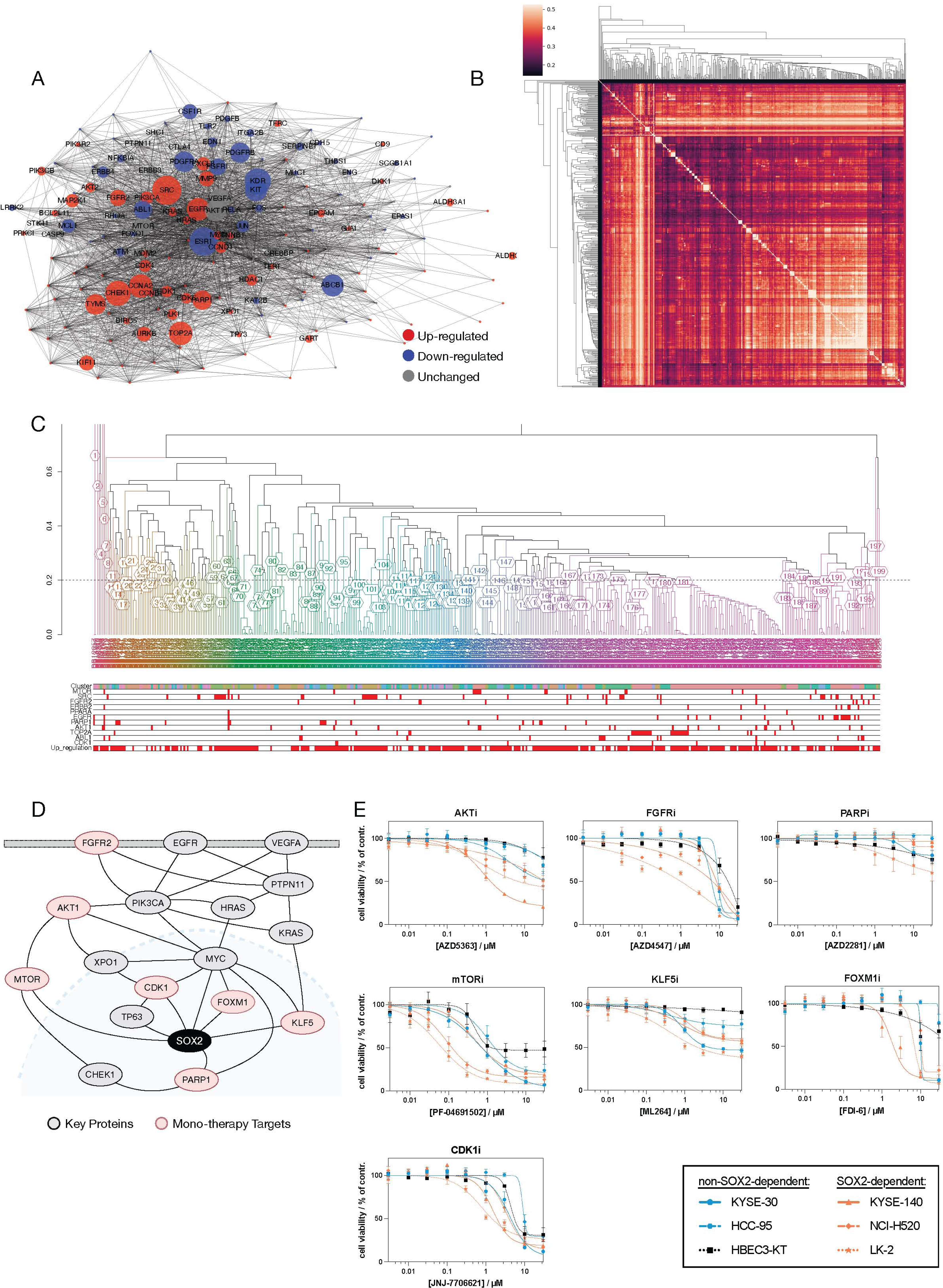
Prioritisation, Drug Simulation, and Experimental Validation of Druggable Monotherapy Targets for SOX2-Dependent LUSC. **A)** Interaction network of candidate targets. Red nodes represent upregulated proteins, and blue nodes represent downregulated proteins in LUSC. Node size corresponds to the number of candidate drugs targeting each protein, with larger nodes such as MTOR and CDK1 indicating highly targeted hubs. **B)** Cluster heatmap of candidate drugs, grouped by chemical structure. Brighter colours indicate higher structural similarity, while white blocks along the diagonal represent clusters of drugs that share structural and functional characteristics. **C)** Hierarchical clustering dendrogram illustrating the relationships between candidate drugs. The colours indicate pathway classifications, such as the PI3K-AKT pathway, cell cycle regulation, and WNT signaling. Upregulated targets are annotated at the bottom, aligning with specific drug clusters. This panel highlights the overrepresentation of upregulated targets in key oncogenic pathways, further supporting their therapeutic potential in SOX2-dependent LUSC. **D)** Druggable pathways associated with the prioritised monotherapy targets. The network highlights connections between FGFR2, PI3K-AKT, and RAS pathways as upstream regulators of SOX2, with implications for therapeutic intervention. **E)** Dose-response curves showing the effect of inhibitors targeting the 7 prioritised monotherapy proteins in SOX2-dependent (orange) and non-SOX2-dependent (blue, black) cell lines. Cells were treated with half-logarithmic concentrations from 3–30,000 nM for 72 hours, and viability was assessed using a resazurin-based assay. Data are presented as mean ± SEM from three independent experiments. The distinct sensitivity of SOX2-dependent cell lines, particularly to inhibitors such as PF-04691502 (MTOR inhibitor) and JNJ-7706621 (CDK1 inhibitor), demonstrates the specificity and efficacy of these prioritised targets.

#### Target Prioritisation

To further prioritise targets, we performed a structural similarity analysis (Figure 4B) to identify clusters of compounds among the 521 drugs, indicating similarity in their target proteins (Supplementary Table 3). Proteins targeted by five or more drugs within each cluster were considered potential primary targets (Figure 5D, Supplementary Table 4). Figure 4C provides a detailed dendrogram showing the hierarchical clustering of drugs and their target proteins. The dendrogram reveals that drugs with similar chemical structures tend to target the same key proteins. At the bottom of the dendrogram, the key druggable targets and all upregulated target proteins are highlighted, providing critical insights into the drug-target landscape. Most of the 521 drugs target proteins that are upregulated in SOX2-dependent LUSC, emphasizing their therapeutic potential in addressing this disease context. The alignment between structurally similar drugs and their shared targets further supports the robustness of the prioritisation approach. Of the 168 key proteins, 84 druggable targets were identified as potential primary targets. Among these, 53 showed efficacy in LUSC, as confirmed by DepMap CRISPR results on LUSC cell lines.

**Figure 5.**
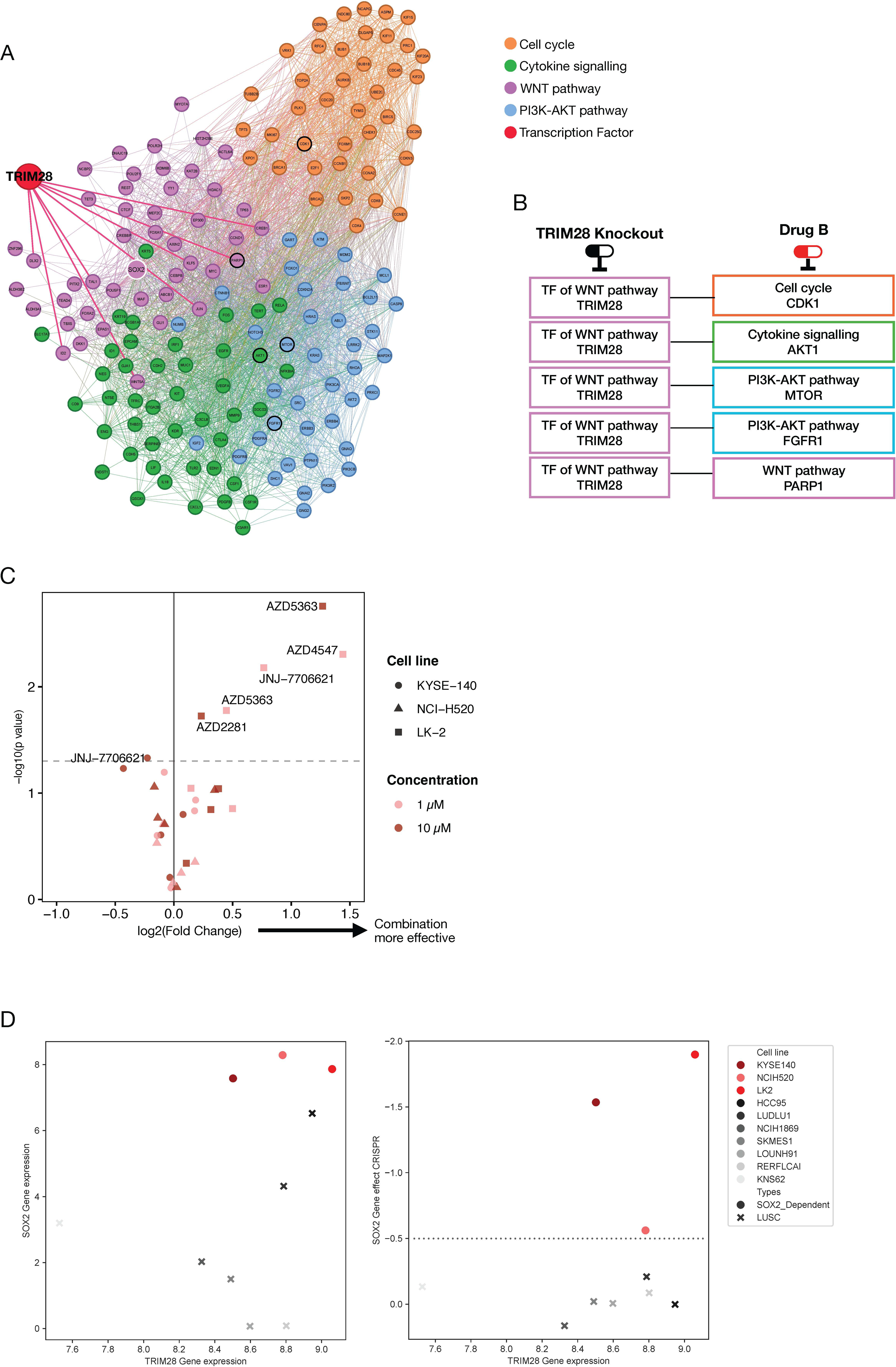
Prioritisation, Network Analysis, and Experimental Validation of Non-Druggable Combination Therapy Targets for SOX2-Dependent LUSC. **A)** TRIM28 was identified as a significantly enriched transcription factor (TF) within the WNT pathway community using a TF enrichment analysis of core network communities. TRIM28 directly regulates eight proteins in the core network, highlighting its critical role in the regulatory landscape of LUSC. **B)** Proposed combination therapy strategies involve TRIM28 knockout paired with drugs targeting proteins from the four communities of the iPANDDA prioritised protein network: cell cycle, cytokine signalling, PI3K-AKT pathway, and WNT pathway. **C)** A volcano plot depicts the effects of drug treatment targeting AKT, CDK1, FGFR, mTOR, and PARP in combination with TRIM28 knockout. Experiments were performed in two SOX2-dependent cell lines (LK2 and KYSE-140). Cells were engineered using guide RNAs (sgRNA) for TRIM28 knockout or a non-targeting control sgRNA, and treatments were applied at 1 µM or 10 µM for 72 hours. Cell viability was assessed, and compounds in the top right quadrant of the plot demonstrate significant enhancement of efficacy in combination with TRIM28 knockout. **D)** The LK-2 cell line exhibited the strongest response to combination therapy (panel C). Elevated expression levels of both TRIM28 and SOX2 in LK-2 cells, along with a pronounced SOX2 gene dependency effect, were observed compared to other LUSC cell lines. This highlights the importance of TRIM28 and SOX2 co-dependency in identifying therapeutic vulnerabilities specific to SOX2- dependent LUSC.

#### Final Prioritisation Using iPANDDA Scoring

The 84 druggable targets were further filtered using the iPANDDA scoring function, which resulted in the selection of seven proteins. Three key criteria were applied during this filtering process:

1. Proteins exhibiting downregulated expression in LUSC were excluded.
2. Proteins encoded by genes with a dependency score greater than 0 in LUSC cell lines (based on DepMap data) were excluded.
3. Only proteins belonging to a chemical structure-based drug cluster comprising at least five drugs were retained as druggable targets.

The remaining proteins were then ranked based on two scores:

1. **Proximity Score (z-score)**: Measures the proximity of a protein to interact with simulated compounds.
2. **Random Walk Restart (RWR) Score**: Quantifies the influence exerted by the SOX2 node on each target protein within the network.

### Pathway Analysis of Druggable Monotherapy Targets

The network depicted in Figure 4D provides a visual representation of the prioritised druggable targets, AKT1, FGFR2, EGFR, VEGFA, PARP1, mTOR, and CDK1, and their connections to SOX2 (Supplementary Table 5). These targets, highlighted as monotherapy candidates, are embedded in the broader network of key proteins influenced by SOX2. The figure emphasizes their central roles in the network and their direct or indirect relationships with SOX2, reinforcing their importance as therapeutic candidates.

From the broader PPI network, we extracted a focused network comprising the 10 prioritised target proteins, including 7 druggable and 3 undruggable targets. Figure 4D highlights two key upstream druggable pathways for SOX2: the PI3K-AKT and FGFR2 pathways. These pathways are fundamental to cancer progression and treatment response. The PI3K-AKT pathway is frequently deregulated or mutated in LUSC, playing a critical role in cell survival and proliferation. Notably, therapeutic targets within this pathway, such as MTOR and CDK1, demonstrate a complex interplay with transcription factors SOX2 and TP63, as depicted in the network. This interplay represents an exploitable vulnerability for therapeutic intervention.

The FGFR2 pathway, another critical axis, is directly associated with SOX2 and represents a promising target for intervention. Both pathways intersect significantly with epithelial-mesenchymal transition (EMT), a key driver of cancer metastasis and recurrence^25,26^. These connections further underscore the strategic importance of targeting these pathways.

This visualisation provides a comprehensive framework for understanding how the prioritised targets align with upstream and downstream regulatory circuits. These insights offer a robust rationale for the selection of these targets for wet-lab validation as monotherapy candidates. Moreover, they lay the groundwork for targeted therapies that can disrupt key regulatory circuits, addressing fundamental mechanisms in SOX2-dependent LUSC pathology.

### Experimental Validation of Prioritised Monotherapy Targets

We considered the identified seven druggable targets of monotherapy for experimental testing for efficacy in SOX2-dependent squamous cell lines. Two of the target proteins, EGFR and VEGFA, from our predicted list have already been tested in clinical trials and were therefore not retested in an in vitro context. EGFR inhibition has been shown to be ineffective in LUSC^27,28^, but targeting VEGF using a monoclonal antibody in combination with PD1 has recently been demonstrated as effective in a landmark study^29^. Notably, prior clinical complications, specifically fatal pulmonary haemorrhage, rather than a lack of efficacy, precluded the use of VEGF-targeting therapies in LUSC^30^. It is noteworthy that iPANDDA identified targets with a sufficient rationale for investment in clinical trials, reaffirming the validity of the computational prioritisation.

For the remaining prioritised targets, AKT1, mTOR, CDK1, FGFR2, and PARP1, and to quantify the efficacy of drugs targeting these proteins, we established a panel of SOX2-dependent and non- SOX2-dependent squamous cell lines (Supplementary Figure 4). We performed a series of dose-response cell population expansion assays using half-logarithmic concentrations ranging from 3 to 30,000 nM for 72 hours.

Importantly, as shown in Figure 4E, there was a clear distinction in the sensitivity of SOX2- dependent cell lines compared to non-SOX2-dependent cell lines when treated with inhibitors targeting AKT1 (AZD5363), mTOR (PF-04691502), and CDK1 (JNJ-7706621). The dose-response curves illustrate a significantly enhanced efficacy of these compounds in SOX2-dependent cell lines, reinforcing their potential as candidate therapeutics for SOX2-dependent LUSC. Figure 4E specifically highlights the steep decline in cell viability for SOX2-dependent cell lines compared to non-SOX2- dependent ones, underscoring the specificity of these inhibitors. This differential response provides robust evidence supporting iPANDDA’s capability to identify novel therapeutic approaches tailored for SOX2-dependent LUSC.

It is important to note that iPANDDA does not imply directionality of the impact of targeting proteins. For instance, while PARP1 was identified as a target protein, both our experimental results (Supplementary Figure 4C) and data from therapeutic screening databases (Supplementary Figure 3C) indicate that PARP inhibition may paradoxically potentiate squamous cancers. This observation aligns with a recent study reporting the failure of PARP inhibition as part of a second-line treatment algorithm for metastatic squamous lung cancer^31^.

### Prioritisation and Validation of Non-Druggable Targets and Combination Therapies

For the remaining 84 proteins not currently targeted by drugs, we performed a similar ranking analysis. For these target proteins lacking known targeting drugs from our simulations, we applied a multi-step filtering process followed by prioritization:

1. Proteins downregulated in lung squamous cell carcinoma (LUSC) samples from the TCGA database were excluded.
2. Proteins with a gene dependency score greater than 0 in LUSC cell lines were filtered out based on data from the DepMap resource.
3. Proteins not directly interacting with the SOX2 transcription factor in our LUSC-specific network (LUSOX) were removed.

After this three-step filtering process, the remaining proteins were prioritized based on their random walk restart (RWR) score from the SOX2 protein node in the network. Three non-druggable target proteins, p63, FOXM1, and KLF5, met the threshold for experimental validation (Figure 5A, Supplementary Table 5, Supplementary Figure 3).

The effective treatment of cancer increasingly involves combination therapies. Leveraging our network analysis, we explored whether novel therapeutic approaches could be developed by combining one of these non-druggable targets with a druggable target. We previously grouped the SOX2-related LUSC pathways into four functional communities: cell cycle, cytokine signalling, PI3K-AKT pathway, and WNT pathway (Figure 3E). Within the WNT pathway community, we identified the transcription factor TRIM28 as a critical regulator, influencing eight proteins that interact with SOX2 (Figure 5A). TRIM28 appears to play a pivotal role in mediating WNT-SOX2 interactions, making it a key candidate for exploring combinatorial targeting strategies. We evaluated the combinatorial efficacy of TRIM28 knockout with inhibition of key druggable proteins from other communities, namely CDK1 (cell cycle), AKT (cytokine signalling), and FGFR (PI3K-AKT pathway) (Figure 5B). This approach aimed to identify synergistic effects across distinct pathways, potentially enhancing treatment efficacy through coordinated mechanisms.

Figure 5C highlights the results of this combination therapy strategy. Notably, in the LK-2 cell line, which exhibits elevated expression of both TRIM28 and SOX2 alongside strong SOX2 dependency, there were significant synergistic responses when TRIM28 knockout was combined with inhibitors of AKT, FGFR, or CDK1. This strong synergy in LK-2 cells underscores the biological relevance of TRIM28 as a potential combination target. Figure 5D provides additional evidence for the importance of TRIM28 in SOX2-dependent LUSC. It shows the relationship between TRIM28 expression and SOX2 expression across cell lines, as well as their dependency scores derived from CRISPR knockout screens. The elevated TRIM28 and SOX2 expression in the LK-2 cell line correlates with the strong combinatorial effect observed in experimental assays, further validating the selection of TRIM28 as a critical node for combination therapies.

TRIM28 is not a currently druggable target, but this work demonstrates that by separating targets into functional communities, iPANDDA can uncover novel approaches to combination therapy. The ability of iPANDDA to recognize and prioritize targets based on their position within distinct network communities, while maintaining focus on SOX2 dependency, highlights its sophisticated understanding of disease-specific biological systems. This capability is particularly critical for LUSC, where the pleiotropic effects of SOX2 suggest that combination therapies may be necessary for effective treatment.

## Discussion

This study introduces iPANDDA, a computational pipeline developed to identify and prioritise therapeutic targets for both monotherapy and combination therapy in SOX2-dependent lung squamous cell carcinoma (LUSC). By integrating and analysing multi-modal datasets, iPANDDA established a robust disease-specific protein-protein interaction (PPI) network, identifying 168 core proteins implicated in LUSC pathogenesis. The stepwise prioritisation of these proteins through drug simulations and network algorithms revealed seven druggable and three non-druggable monotherapy candidates (Figure 6).

**Figure 6.**
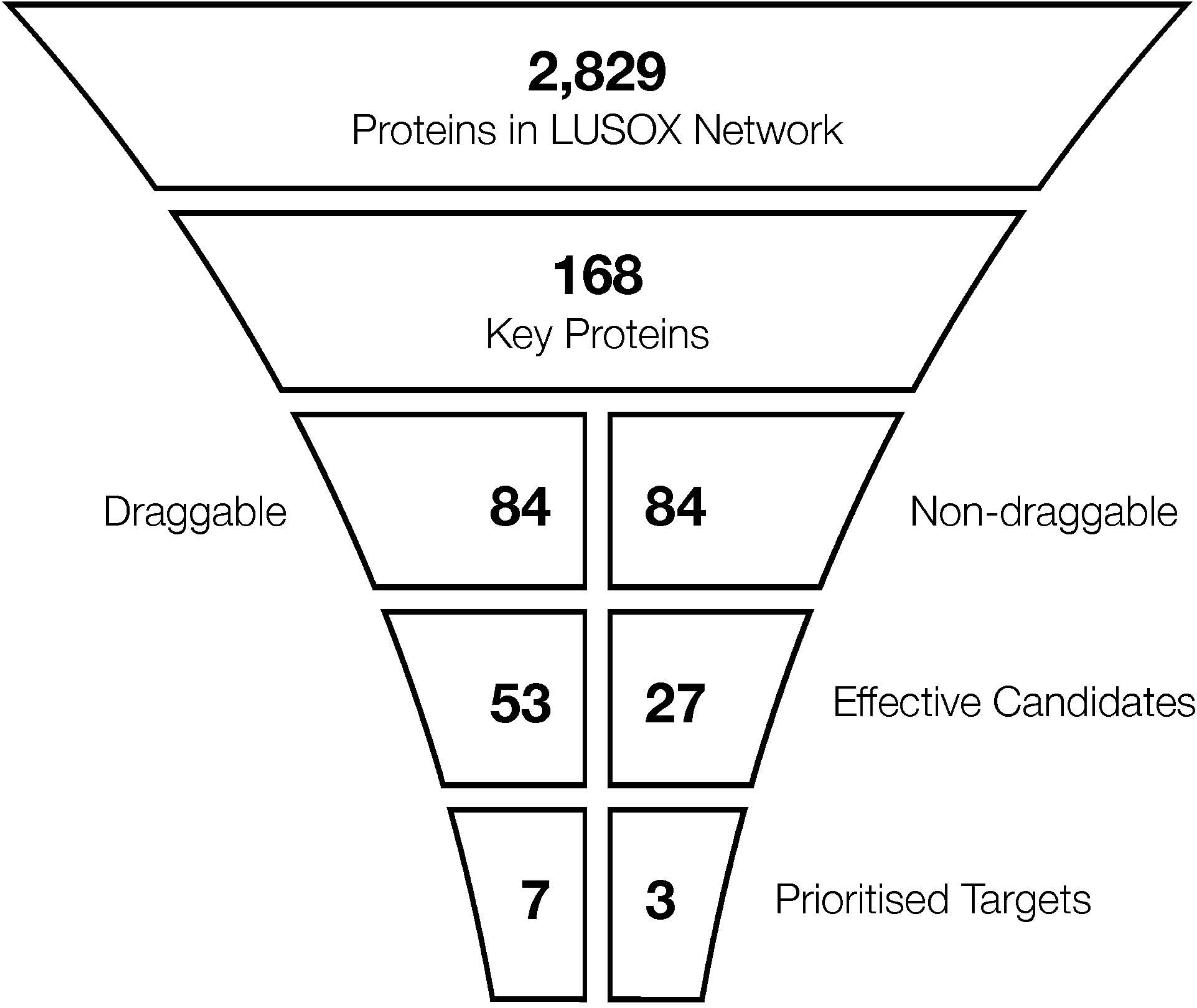
Stepwise Prioritisation of Druggable and Non-Druggable Targets in SOX2-Dependent LUSC. A systematic pipeline from the LUSOX network (2,829 proteins) identified 168 key proteins through network analysis. Drug simulations stratified these into druggable (n = 84) and non-druggable (n = 84) proteins. Parallel prioritisation approaches refined druggable proteins to 53 effective candidates and non-druggable proteins to 27, further narrowed to seven druggable and three non-druggable prioritised targets for monotherapy.

Figure 6 summarises the stepwise prioritisation process, starting with 2,829 proteins in the LUSOX network. Through systematic network analysis, 168 key proteins were identified based on centrality metrics and biological relevance. Drug simulations were then performed using 6,009 compounds from DrugBank to evaluate their potential interactions with the 168 key proteins, resulting in 521 candidate drugs targeting these proteins. Based on the simulation results, the 168 key proteins were stratified into two groups: druggable (n = 84) and non-druggable (n = 84).

From this point, a parallel approach was taken to further refine the druggable and non-druggable groups. For the druggable proteins, dependency scores, expression levels, and clustering within drug-target interaction networks reduced the pool from 84 to 53 proteins. Simultaneously, the candidate drugs were refined from 521 to 217 with potential LUSC therapeutic relevance. Among these, 114 drugs overlapped with cancer drugs, underscoring their biological and therapeutic significance. For the non-druggable proteins, similar dependency scores and network analyses reduced the pool to 27 effective candidates.

Further prioritisation narrowed the druggable proteins to seven high-priority targets based on expert reviews, experimental feasibility, and clustering analyses. In parallel, the non-druggable proteins were refined to three prioritised targets using combinatorial therapy assessments (Figure 5A and 5B), alongside expert reviews and feasibility considerations. Experimental validation confirmed the efficacy of inhibitors targeting AKT1, MTOR, and CDK1 for monotherapy in SOX2-dependent LUSC cell lines, while combinatorial evaluations demonstrated the relevance of non-druggable targets for combination therapies. These results highlight the systematic and iterative nature of iPANDDA in narrowing down a large protein network to actionable targets and therapeutic candidates.

In addition to monotherapy targets, iPANDDA identified TRIM28 as a critical regulator within the SOX2-centric network. TRIM28 emerged as a promising combination therapy target, particularly when paired with inhibitors of AKT, CDK1, or FGFR1 from other core pathway communities. Experimental evaluations demonstrated selective effectiveness of these combinations in SOX2- dependent LUSC models, particularly in the LK-2 cell line, which exhibits high SOX2 and TRIM28 dependency. These findings highlight iPANDDA’s capacity to address the molecular complexities of LUSC and identify actionable opportunities for combination therapies.

The integration of diverse datasets and systematic prioritisation provided a comprehensive framework for identifying both well-characterised and novel targets. This approach significantly expands the therapeutic landscape for SOX2-dependent LUSC. The in vitro validation experiments further emphasised the importance of tumour molecular characteristics—specifically SOX2 dependency—in determining therapeutic response. The identification of TRIM28 as a combination therapy target illustrates the transferability of iPANDDA’s methodology, offering a template for similar analyses in other cancers.

However, certain limitations should be acknowledged. The predictive power of iPANDDA relies on the completeness and accuracy of protein interaction and drug target databases. Additionally, current in vitro models may not fully replicate the complexity and heterogeneity of the human tumour microenvironment, necessitating complementary in vivo studies to validate findings further.

Despite these limitations, iPANDDA represents an innovative and comprehensive approach that integrates multi-modal datasets within a unified analytical framework, offering a systematic solution to therapeutic target discovery. While the current study focused on SOX2-dependent LUSC, this methodology is broadly applicable to other cancers, enabling the identification of novel targets across a wide range of disease contexts. Future efforts will aim to incorporate machine learning techniques to refine target prioritisation, particularly for mono- and combination therapies, further enhancing the pipeline’s accuracy and translational potential.

---

**Supplementary Figure 1: Principal Component Analysis of Disease and Non-Disease Samples in TCGA and GTEx**

Principal Component Analysis (PCA) of RNA-seq data from disease (TCGA) and non-disease samples (TCGA and GTEx). Each point represents a sample, with color denoting the sample type: disease samples from TCGA (red), non-disease samples from TCGA (blue), and non-disease samples from GTEx (green). PCA was performed on normalized expression profiles, illustrating the separation between disease and non-disease states.

**Supplementary Figure 2: Circos plots of LUSOX**

**A)** Highlight of interactions from SOX2 PPI

**B)** Highlight of interactions from Master proteins of LUSC

**C)** Highlight of interactions from LUSC drugs

**D)** Highlight of interactions from TF signatures

**Supplementary Figure 3: Targets for monotherapy**

**A)** Relationship between SOX2 and prioritised targets Pearson correlation coefficient R for target gene vs. SOX2 mRNA expression with significance of correlation. Data from the TCGA LUSC project. Higher value (red) means higher correlation of target expression with SOX2 expression. mRNA expression fold change in KYSE-140 cells after SOX2 knockout. Significance from t-test comparing SOX2 knockout with control. Data from Liu et al. Higher value (red) means overexpressed after SOX2 knockout. Regulatory potential score for the SOX2 TF to regulate the target genes. Higher score (red) means higher regulatory potential of target gene by SOX2.

**B)** Expression of prioritised targets in squamous tumours, LUAD and normal lung tissue in TCGA datasets.

**C)** The correlation between SOX2 dependency and drug sensitivity, measured by area under the curve (AUC), was analysed for drugs targeting the prioritised targets identified from the iPANDDA pipeline. This analysis utilised drug sensitivity data from the Genomics of Drug Sensitivity in Cancer (GDSC) project across 77 squamous cancer cell lines. Pearson correlation coefficients between SOX2 dependency and drug sensitivity AUC values were calculated, along with their statistical significance.

**D)** Dependency of target candidates in cell lines: Density plots showing the number of cell lines with target gene dependency probabilities. Higher gene dependency probability value means the gene is more essential in a cell line. All screened cell lines (n = 1086) are shown in red, squamous cell lines (n = 119) are shown in blue, and LUSC cell lines (n = 22) are highlighted with green stripes.

**Supplementary Figure 4: Target validation for monotherapy**

**A)** Characterisation of cell lines with SOX2 copy number (CN) ratio (1 means same CN as ploidy), SOX2 dependency probability (higher means more dependent), and mRNA expression of squamous markers. Data from the CCLE and DepMap projects.

**B)** Western blot showing the expression of selected squamous markers in the cell line panel.

**C)** Bar charts showing the mRNA expression of target genes in the cell line panel by RT-qPCR. Gene expression was normalised to TBP and is shown relative to HBEC3-KT. Data presented as mean ± SEM of N = 3 biological replicates (n = 3 technical replicates).

**D)** Heatmap showing drug sensitivity in SOX2-dependent and non-SOX2-dependent cell lines by area under the curve (AUC) values calculated from dose-response curves (see Figure 3E).

**Supplementary Figure 5: Target validation for combination therapy**

**A)** Schematic overview of the experimental design to evaluate combination therapies.

**B)** Individual plots of cell viability after treatment of cell lines with selected compounds in combination with sgNTC or sgTRIM28. CRISPR/Cas9-engineered cells were expanded and then treated with 1 µM or 10 µM of selected compounds for 72h and cell viability assessed. Data presented as means ± SEM of N = 3 biological replicates (n = 3) technical replicates.

**Supplementary Table 1**: Multi-modal data integration result

**Supplementary Table 2**: Network analysis result

**Supplementary Table 3**: Drug simulation result

**Supplementary Table 4**: Drug cluster table

**Supplementary Table 5**: Target prioritisation result

## Methods

### Chromatin immunoprecipitation (ChIP)

Cells cultured in OTC and untreated/treated with 2μg/ml Dox for 4 days were harvested. To harvest cells cultured in the OTC, medium was removed from both lower and upper chamber and the porous membrane of each transwell was wrapped in cling film to ensure the cells were fully submerged into the solutions. The collagen disks were washed twice with 500μL of Phosphate-Buffered Saline (PBS- Sigma) at RT and trypsinised with 3x trypsin-EDTA for 12min at 37°C. Trypsinised cells from 3 transwells (under the same experimental condition) were collected into a microcentrifuge tubes containing DMEM/FBS and centrifuged at 1100rpm. Approximately 1×106 cells were used for ChIP assay, carried out according to the manufacturer’s instructions (Magna ChIPTM G, Millipore, Cat No. 17-611). Briefly, cells from 5 different transwells were collected into the same tube and cross-linked by resuspending the cell pellet in 10ml of KSFM+supplements with 275μL of 37% formaldehyde for 10min at RT. The reaction was quenched by addition of 1ml of 10x glycine and washed twice with cold 1x PBS. The fixed cells were resuspended in cell lysis buffer. Nuclei were collected by centrifugation at 2000rcf, and resuspended in nuclear lysis buffer. Samples were then sonicated to produce chromatin fragments of an average of 500 bp. Chromatin shearing was performed in 130μL microtubes (with 130μL total volume) using the sonicator Covaris S2 series and the following programme: Duty cycle: 2%. Intensity: 3. Cycles per burst: 200. Duration: 12min.

50μL of sheared chromatin was incubated with 20μL protein G magnetic beads 5 µg of either Goat anti- SOX2 antibody (AF2018) or Goat anti-IgG control antibody (AB-108) overnight at 4°C with rotation. Next day, the protein–DNA complex was washed, the cross-link was reversed and proteins were removed by proteinase K treatment. DNA was purified using spin columns and finally eluted in 50μL of elution buffer.

For the validation of immunoprecipitation, the sheared chromatin was incubated with a validated antibody and a negative control followed by PCR of the eluted DNA (ChIPAb+ RNA Pol II – ChIP Validated Antibody and Primer set, Millipore, Cat No. 17-672).

Initial sample quality control of pre-fragmented DNA was performed using a Tapestation DNA 1000 High sensitivity Screen tape (Agilent, Cheadle UK). Sequencing ready libraries were prepared from samples using the Hyper Prep DNA Library preparation kit (Kapa Biosystems, London UK) and indexed for pooling using NextFlex DNA barcoded adapters (Bioo Scientific, Austin TX US). Libraries were quantified on a Tapestation DNA 1000 Screen tape and by qPCR using an NGS Library Quantification Kit (KAPA Biosystems) on an AriaMx qPCR system (Agilent). Libraries were then normalised, pooled, diluted and denatured for sequencing on the NextSeq 500 (Illumina, Chesterford UK). Sequencing was performed using a high output flow cell with 2×75 cycles of sequencing.

### ChIP-Seq Data Processing

Raw sequencing reads were assessed for quality using FastQC. High-quality reads were aligned to the human reference genome (GRCh38) using Bowtie2, and alignment statistics were analyzed using Samtools. Aligned BAM files were sorted and indexed with Samtools, and duplicate reads were marked and removed using Picard Tools to minimize artifacts. Normalized coverage tracks (bigWig files) were generated using bamCoverage from the deepTools suite to visualize genome-wide SOX2 binding profiles. Signal differences between ChIP samples and controls were quantified using bamCompare from deepTools, enabling the generation of differential signal profiles.

### Multi-modal signatures

Transcriptomics disease signatures (TCGA & GTEx): Lung squamous cell carcinoma (LUSC)-specific cancer genes were identified by analyzing transcript per million (TPM) values from the LUSC-TCGA TARGET GTEx dataset, a combined cohort of TCGA, TARGET, and GTEx data provided by UCSC Xena^30^/01/2025 09:29:00. The full dataset comprises 17,221 samples, including 9,807 from TCGA and 7,414 from GTEx. For LUSC, the data comprise 498 disease samples (TCGA) and 337 healthy samples, including 50 from TCGA and 287 from GTEx. For each disease sample, the fold-change was calculated as the ratio to the average of healthy samples (TCGA + GTEx). Subsequently, genes were filtered using an ANOVA test to identify those exhibiting significant differential expression in LUSC compared to other cancer types in the TCGA pan-cancer dataset (31 cancer types) and matched normal tissues from the GTEx dataset (30 tissue types). The genes were ranked based on their ANOVA p-values from these comparisons. A distinct sharp increase in p-values was observed after the threshold of 1e-150. Therefore, genes with p-values ≤ 1e-150 were selected as the LUSC-specific gene set, applying a highly stringent cutoff to ensure high specificity.

Using this LUSC-specific gene set, hierarchical clustering was performed on samples from 31 cancer types present in the TCGA database to explore cancer type-specific expression patterns and potential subgroups or subtypes based on the expression profiles of these genes.

LUSC functional signatures: LUSC-associated genes were selected from the Open Targets 19.09^14^. The Open Targets platform integrates evidence from various data sources, including genetics, genomics, transcriptomics, drug information, animal models, and scientific literature, to compute association scores between targets and diseases. Genes with an overall association score of >= 0.8 were considered as LUSC-associated genes. We chose a threshold value of 0.8 for inclusion based on a disease enrichment analysis (Enrichr^32^) performed on gene lists with a range of overall association score thresholds (0.75, 0.8, 0.85, 0.9). The threshold of 0.8 exhibited the lowest p-value for enrichment in squamous cell carcinoma of lung, indicating the most significant association with the disease of interest.

Drug Target Signatures: Drug target signatures consist of proteins that are established as targets of drugs currently in Phase 3 or higher clinical trials for lung squamous cell carcinoma (LUSC). The data on LUSC drugs and their corresponding protein targets were sourced from the Open Targets 19.09 database.

### LUSOX network construction

The LUSOX network was constructed using a multi-modal signature PPI network from the STRING database. Only interactions with a confidence score of greater than medium (0.4) were used. Networks were visualized using Gephi 0.9.2^33^ (Figure 2C).

### LUSOX network community detection

The Louvain community detection algorithm, implemented in the Gephi software, was applied to LUSOX to identify communities. The Louvain method is a hierarchical clustering approach that optimizes modularity to detect communities of nodes that are more densely connected within themselves compared to connections between communities.

### Network analysis to identify key proteins

Eigenvector centrality, degree centrality, betweenness centrality and random walk with restart (RWR) were utilized to identify key proteins in the LUSOX network. The LUSOX network was represented by an adjacency matrix *A*, where *A_ij_*= 1 there is an edge between nodes *i* and *j* or *A_ij_*= 0 otherwise. The eigenvector centrality *x_i_* was defined as

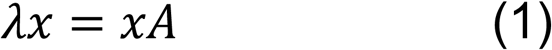

where *x* is an eigenvector of the adjacency matrix *A* with eigenvalue *λ*. If *λ* is the largest eigenvalue of the adjacency matrix *A*, there is a unique solution *x*, all centrality values are positive ^34^. Degree centrality of node *i* was defined as

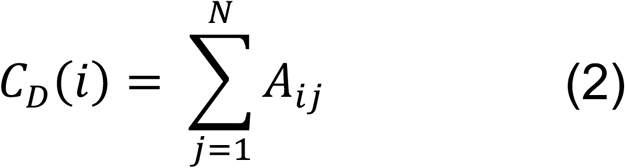

where *N* is the number of nodes in the LUSOX network. Betweenness centrality of a node *i* was defined as

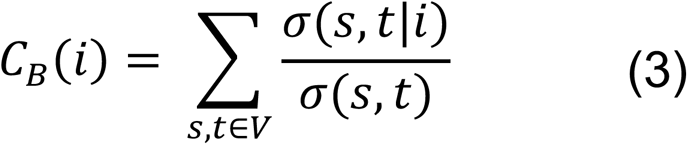

where *V* is the set of nodes, *σ*(*s*, *t*) is the total number of shortest paths between *s* and *t*, and *σ*(*s*, *t*|*i*) is the number of shortest paths between *s* and *t* paths passing through node *i*. If *s* = *t*, *σ*(*s*, *t*) = 1, and if *i* ∈ *s*, *t*, *σ*(*s*, *t*|*i*) = 0.

Eigenvector centrality was used to identify the most influential proteins in the network. If a protein frequently interacts with other proteins that also have high eigenvector centrality, then the protein will have high eigenvector centrality. Degree centrality was used to identify the hub proteins in the network. Betweenness centrality was used to identify the bottleneck proteins in the network. A RWR algorithm was used to see which human proteins are most affected by SOX2 protein. To do this, we used SOX2 as the starting point of RWR. The RWR parameters were (1) a restart probability of 0.15, (2) a maximum iteration number of 100, and (3) an error tolerance of 1e-06. We assigned edge betweenness centrality as an edge score on the LUSOX. The RWR calculated a score per protein in the LUSOX network, which indicates how much of a given protein was influenced by SOX2. The algorithms were implemented in the Python package NetworkX (v 2.2) ^35^.

Permutation tests were performed 1,000 times to identify significant proteins for each of the network centrality algorithms. In 1,000 permutation tests, each test generated a random network with a preserved degree distribution of the original network, the LUSOX network. To generate a random network, we reconnected the edge in the LUSOX network and swiped the node. So, the random network in each permutation test has at least 66% of the rewired edges. Then, in the permutation test, we applied the network algorithm and obtained the cumulative results of the network algorithm. These cumulative results were used to calculate the empirical p-value of the network algorithm. We combined the four permutation test results to determine the final set of key proteins that have an empirical p-value <=0.01 in either result.

### Drug-Target interactions

Approved drugs were collected from ChEMBL^36^ and DrugBank^37^. Drug-target interaction information was collected from DrugBank(v 5.1)^37^, STITCH (confidence score > 0.9)^38^ and Cheng, et al^39^.

### Drug simulation

Approved drugs were collected from ChEMBL ^36^ and DrugBank ^37^. Drug-target interaction information was collected from DrugBank (v 5.1) ^37^, STITCH (v 5.0,confidence score > 0.9) ^38^ and Cheng, *et al.* ^39^. *in-silico* network-based proximity analysis was conducted for key proteins from the LUSOX network. Given K, the set of key proteins from LUSOX networks, and T, the set of drug targets, the network proximity(equation (5)) of K with the target set of T of each approved drug where d(k, t) the shortest path length between nodes k ∈ K and t ∈ T in the human PPIs ^39^ was executed. Closest distance measure was used to calculate the distance between a given drug’s targets and key proteins in the LUSOX network, because it previously showed best performance in drug-disease pair prediction in the study of Guney *et al.,* ^40^.

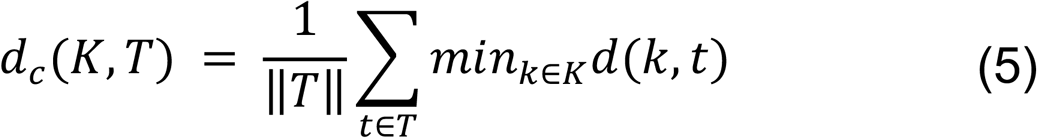

To assess the significance of the distance between a key protein in the LUSOX network and a drug *d_c_*(*K*, *T*), the distance was converted to Z-score based on permutation tests by using

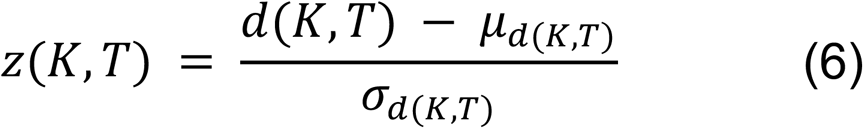

The permutation tests were repeated 1,000 times, each time with two randomly selected gene sets. There are few high degree nodes due to the scale-free network of the human protein–protein interaction network. To avoid repetitive selection of the same high degree nodes during random selection, we used a binning approach with at least 100 nodes in a bin. In the binning approach, nodes in same bin have a similar node degree to maintain node degree distribution for random selection. When we randomly select a set of genes, we perform a random selection among proteins from all bins to verify that the minimum node degree was less than the minimum node degree of the selected gene set and the maximum node degree was greater than the maximum node degree of the selected gene set. The corresponding p-value was calculated based on the permutation test results. Drug to SOX2-dependent LUSC associations with a Z-score of less than −2 were considered significantly proximal ^40^.

### Prioritisation of key proteins for therapy

Three key criteria are employed to filter potential protein targets: 1) Proteins exhibiting downregulated expression in lung squamous cell carcinoma (LUSC) samples from the TCGA database are excluded. 2) Proteins encoded by genes with a dependency score greater than 0 in LUSC cell lines, based on data from the DepMap database^16^, are filtered out. 3) Only proteins belonging to a chemical structure-based drug cluster comprising at least five drugs are considered as druggable targets.

The remaining proteins after this triple-filtering process are then prioritized based on two scores:

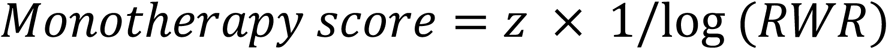

i) Proximity score (z-score) from drug simulations

ii) Random walk restart (RWR) score from the SOX2 protein

The proximity score calculated the proximity of a protein to interact with the compounds simulated, while the RWR score quantifies the influence exerted by the SOX2 node on each target protein within a network analysis framework. Thus, targets are ranked according to their predicted druggability (proximity score) and functional relationship with the SOX2 TF (RWR score).

For target proteins lacking known targeting drugs from our simulations, we applied a different multi-step filtering process followed by prioritization: 1) Proteins downregulated in lung squamous cell carcinoma (LUSC) samples from the TCGA database were excluded. 2) Proteins with a gene dependency score greater than 0 in LUSC cell lines were filtered out based on data from the DepMap resource. 3) Proteins not directly interacting with the SOX2 TF in our LUSC-specific network (LUSOX) were removed.

After this three-step filtering process, the remaining proteins were prioritized based on their random walk restart (RWR) score from the SOX2 protein node in the network.

### Key protein network community detection

Key protein network is derived from the LUSOX network, which consists of 168 proteins. The Louvain community detection algorithm, implemented in the Gephi software, was applied to this key protein network to identify communise.

### Identification of combination therapy

Combination drug targets were selected based on the following criteria: 1) The pair of targets was chosen such that each drug targets a different community. 2) One target was selected as the TF (TRIM28) that regulates the entire community containing SOX2. 3) The other target was selected from the list of monotherapy candidates, with each candidate belonging to a different community within the network.

TF enrichment analysis was performed on the community/module containing the SOX2 protein, identified as the WNT pathway community. The enrichment was evaluated using the Enricr and the ENCODE TF binding site dataset. An over-representation test was conducted, and TRIM28 was identified as this community’s top enriched TF.

### Cell viability assay

Cells were incubated for 24h before adding drugs in DMSO in half-logarithmic concentrations from 3- 30,000 nM. After incubation for 72h, the alamarBlue assay was used to assess cell viability by measuring fluorescence intensity. Dose-response curves were fitted in Graphpad Prism using data points from at least three biological replicates.

### In-vitro validation of combination therapies

PX458 was a gift from Feng Zhang (Addgene # 48138). Previously validated sgRNA sequences targeting TRIM28 and a non-targeting control were cloned into PX458 to express sgRNAs and Cas9, and transfected into cell lines using lipofectamine LTX. After 72h, GFP-expressing cells were sorted on the Aria III cell sorter, expanded and used in cell viability assays to assess combinatorial effects.

### Phenotyping of cell line markers

Cells were harvested and protein lysates immunoblotted as described in Correia et al., 2017 ^41^. RNA was harvested using the RNeasy Mini kit, cDNA synthesised with the High Capacity cDNA Reverse Transcription kit, and RT-qPCR performed with SYBR Green reagents. Data was analysed using the ΔΔCt method and were normalised to TBP expression.

## Funding

W.H. M.M. and N.H. were funded by LifeArc. D.K. was supported by the MRC DTP at the University of Cambridge. L.C. and F.M. were funded by the Wellcome Trust (WT097143MA). F.M. and W.G. were supported by the BBSRC (BB/W014564/1) and the NC3Rs (NC/S001204/1). This research was supported by the National Institute for Health and Care Research (NIHR) Cambridge Biomedical Research Centre (NIHR203312*). The views expressed are those of the authors and not necessarily those of the NIHR or the Department of Health and Social Care.

## Data and materials availability

All data needed to evaluate the conclusions in the paper are present in the paper and/or the Supplementary Materials. The code and input data that were used for this study are available at TBA. Additional data related to this paper may be requested from the authors.

## Competing interests

N.H. is a cofounder of KURE.ai and CardiaTec Biosciences. W.H and M.M. are employees of CardiaTec Biosciences. D.K. is an employee of The Boston Consulting Group, Berlin, Germany. L.C. is an employee of Adaptimmune. R.H. is an employee of Cancer Research Horizons. All other authors declare that they have no competing interests.

## Author contributions

Conceptualization: W.H., D.K. F.M., and N.H.

Methodology: W.H., D.K. F.M., and N.H.

Investigation: W.H., D.K. W.G., L.C.

Visualization: W.H., D.K. F.M., and N.H.

Supervision: F.M. and N.H.

Writing—original draft: W.H. D.K., M.M. F.M., and N.H.

Writing—review & editing: W.H. D.K., W.G., M.M. R.H. F.M., and N.H.

