## Supplementary Table 5 for "Utilising multi-modal data-driven network analysis to identify monotherapy and combinational therapy targets in SOX2-dependent squamous cell lung cancer"

| Target | Description | Implication in LUSC/SQC | Link with SOX2 |
| --- | --- | --- | --- |
| Currently druggable targets (“known targets”) | | |  |
| **AKT1** | AKT serine/threonine kinase 1 | ·   Gene amplified in 4% of LUSC patients^1^ | ·   SOX2 promotes ESCC growth via AKT/mTOR^2^ |
|  |  | ·   PI3K/AKT/mTOR overactive in LUSC^3^ | ·   AKT stabilises SOX2^4^ |
|  |  | ·   MK-2206 promising results^5^ |  |
| **FGFR2** | Fibroblast growth factor receptor 2 | ·   Mutated in 3% of LUSC patients^6^ | ·   Cooperates with SOX2 to transform tracheobronchial epithelial cells^7^ |
|  |  | ·   *FGFR1* amplifications and fusions more common^8^ | ·   Required for maintenance of stemness via SOX2^9^ |
|  |  |  | ·   AZD4547 reduces SOX2 expression^10^ |
| **EGFR** | Epidermal growth factor receptor | ·   Gene amplified in 8% of LUSC patients^1^ | ·   *EGFR* is SOX2 TF target^11^ |
|  |  | ·   Overexpression and amplification more common in LUSC than LUAD^12,13^ | ·   EGFR signalling activates SOX2^14^ |
| **VEGFA** | Vascular endothelial growth factor A | ·   Angiogenesis as hallmark of cancer^15^ | ·   Activates SOX2 to promote EMT^16^ |
|  |  | ·   Ramucirumab (VEGFR) approved for LUSC^17^ |  |
| **PARP1** | Poly(ADP-ribose) polymerase 1 | ·   High mutational burden and genomic instability^1^ | ·   SOX2 TF requires PARP1 to bind certain motifs^18^ |
|  |  | ·   Olaparib showed improved OS in LUSC^19^ |  |
| **mTOR** | Mechanistic target of rapamycin kinase | ·   PI3K/AKT/mTOR overactive in LUSC^3^ | ·   SOX2 promotes ESCC growth via AKT/mTOR^2^ |
|  |  | ·   AZD2014 promising results but discontinued^20^ | ·   mTOR inhibition reduces SOX2 expression^21^ |
| **CDK1** | Cyclin dependent kinase 1 | ·   Sustained cell cycle through deregulation of cell cycle regulators^22^ | ·   Direct interaction with SOX2^23^ |
|  |  |  | ·   Promotes stemness via SOX2^24^ |
| Currently not druggable targets (“novel targets”) | | |  |
| **p63** | Tumour protein p63 | ·   Gene amplified in 50% of LUSC patients^1^ | ·   Co-amplified with SOX2 on chromosome 3q^25^ |
|  |  | ·   Tumour promoting ΔNp63 is dominant isoform in SQC^26^ | ·   Cooperates with SOX2 to induce EGFR^27^ |
| **FOXM1** | Forkhead box M1 | ·   Overexpression promotes EMT^28^ | ·   Binds to *SOX2* promoter^29^ |
|  |  | ·   Marker for bad prognosis^30^ | ·   Downregulation reduces SOX2 expression^29^ |
|  |  |  | ·   SOX2 activates FOXM1^31^ |
| **KLF5** | KLF transcription factor 5 | ·   Induces EGFR expression^32^ | ·   Interacts with SOX2 and p63^33^ |
|  |  | ·   Can act as oncogene or tumour suppressor depending on cell context^34^ | ·   Downregulation reduces SOX2 expression and vice versa^35^ |

Supplementary table 6 Target prioritisation result
