## Supplementary Figures for "Utilising multi-modal data-driven network analysis to identify monotherapy and combinational therapy targets in SOX2-dependent squamous cell lung cancer"

Supplementary Figure 1

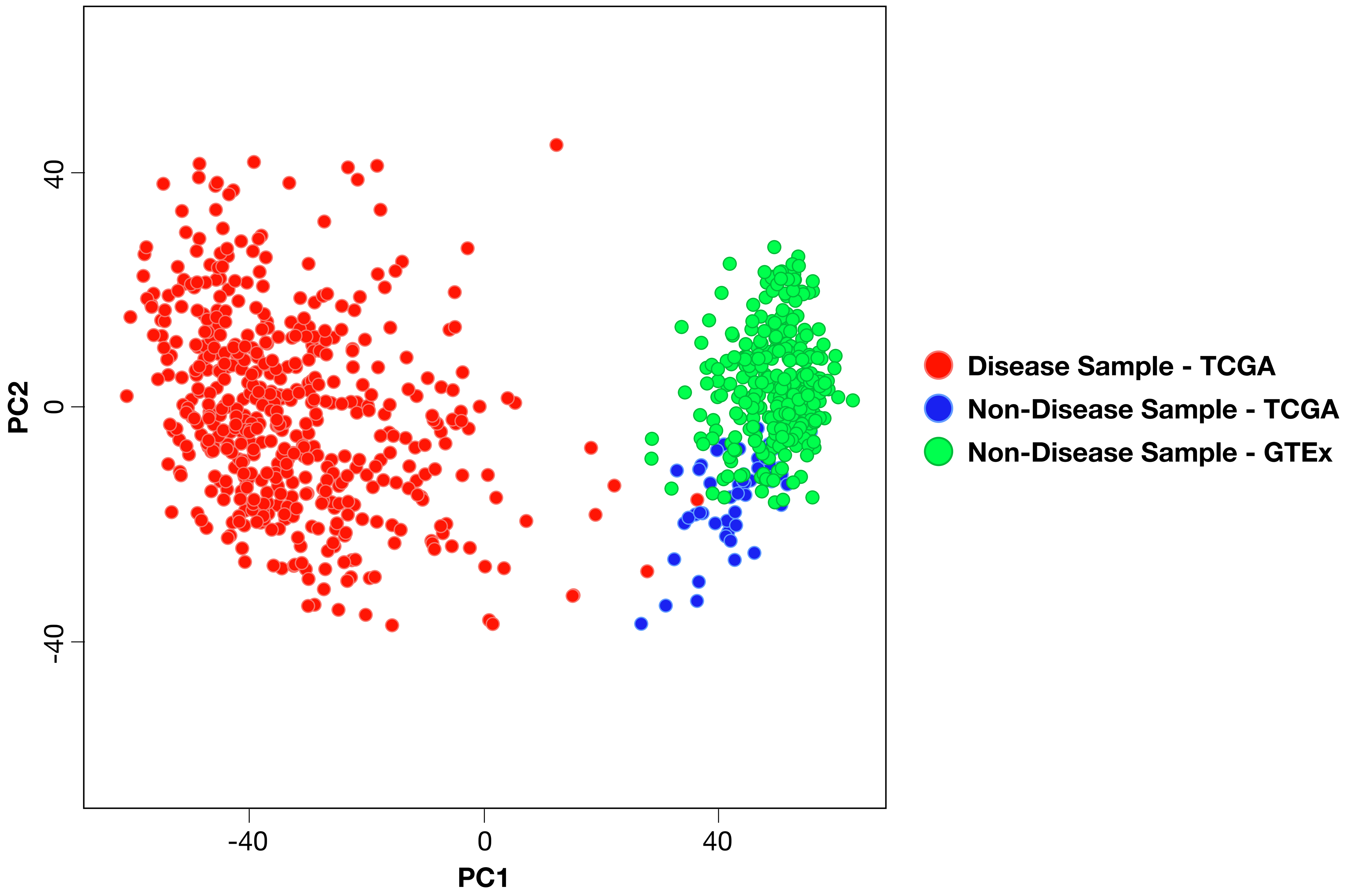

Supplementary Figure 2

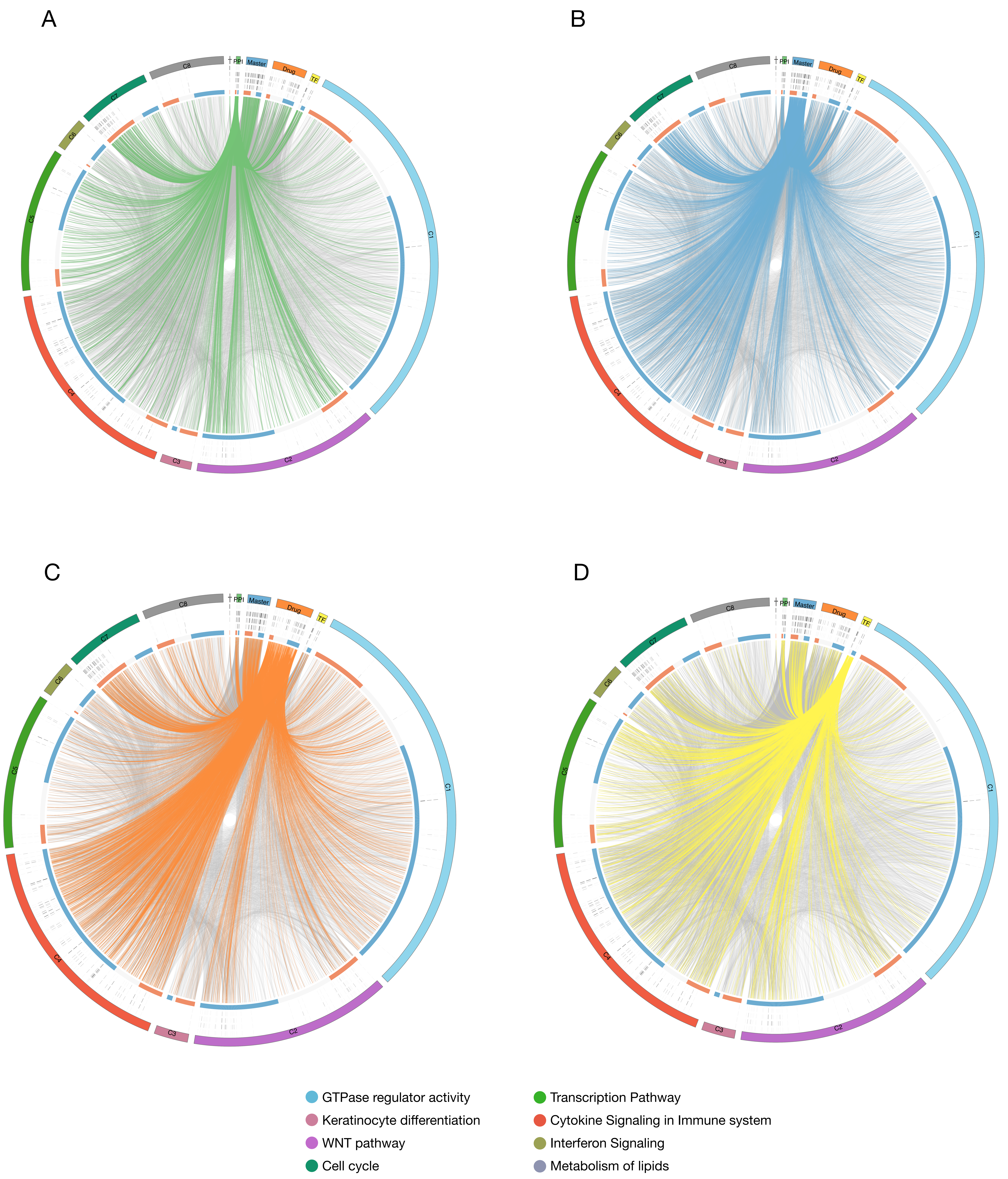

### Supplementary Figure 3

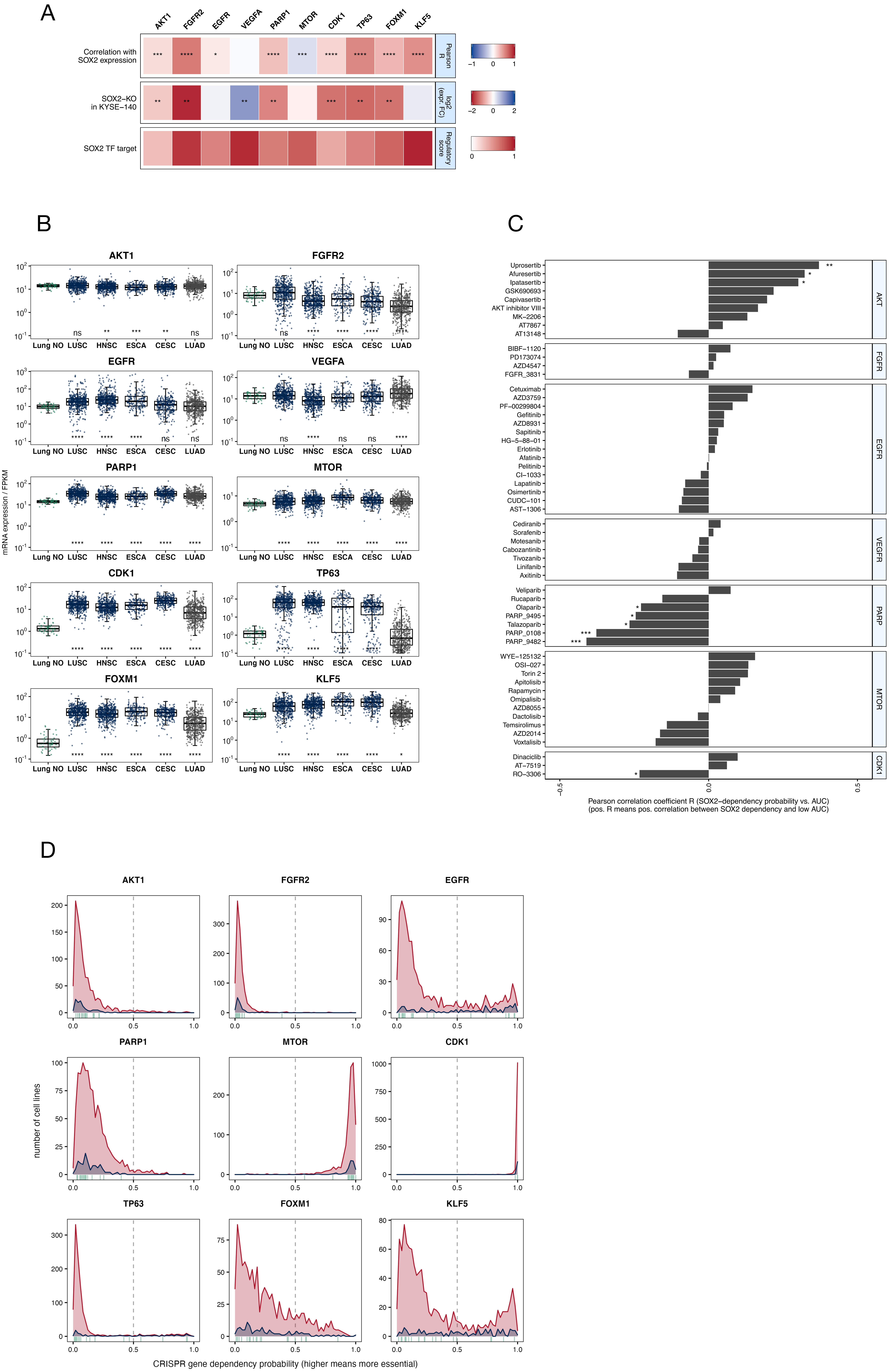

Supplementary Figure 4

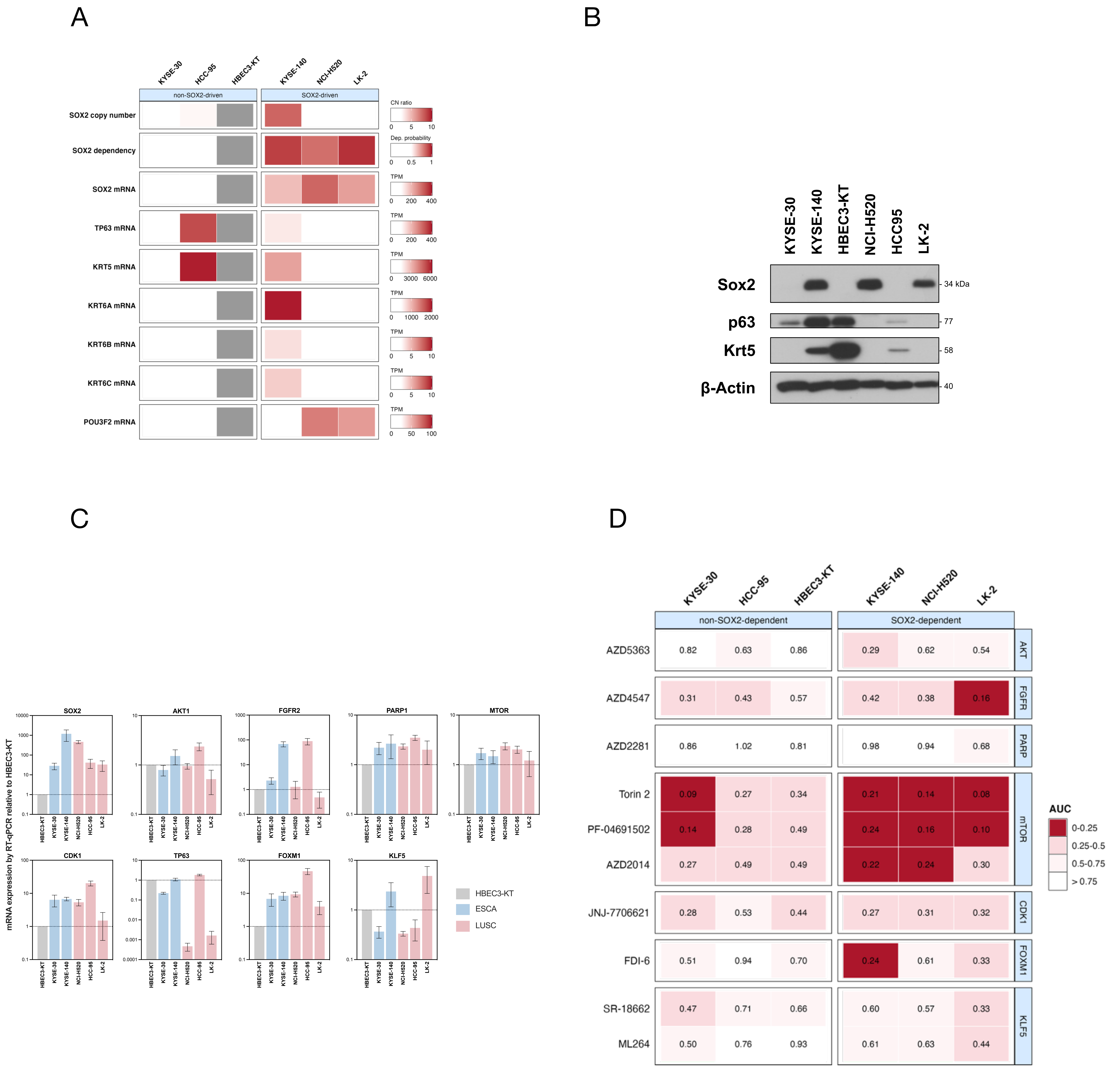

Supplementary Figure 5

A

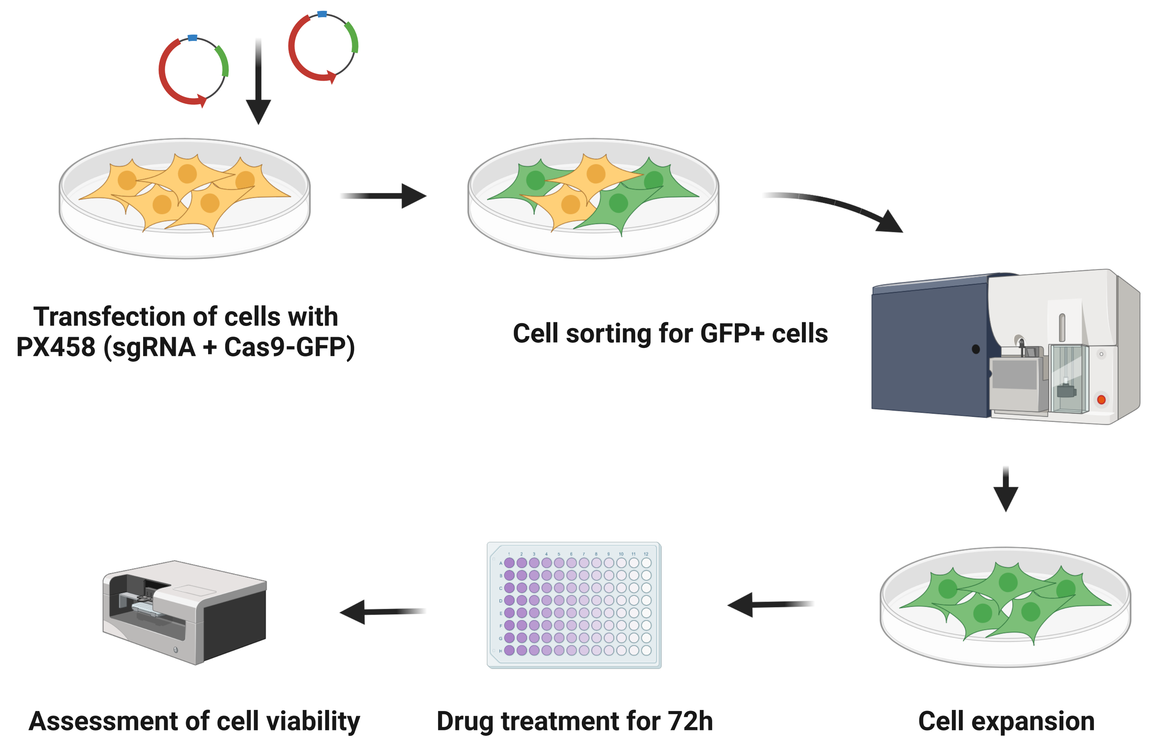


B

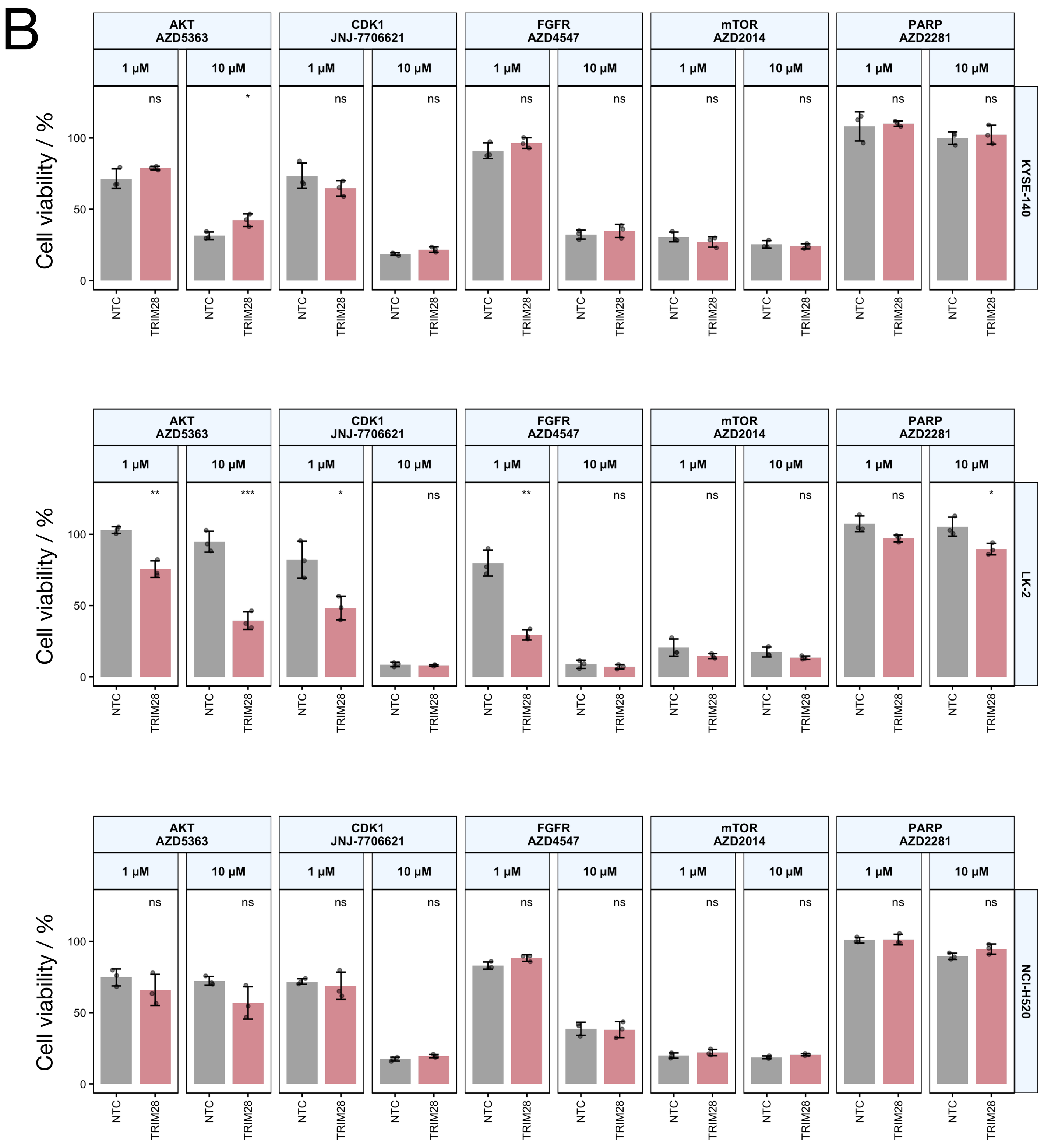
